# Critical assessment of *E. coli* genome-scale metabolic model with high-throughput mutant fitness data

**DOI:** 10.1101/2023.01.05.522875

**Authors:** David B. Bernstein, Batu Akkas, Morgan N. Price, Adam P. Arkin

**Affiliations:** Department of Bioengineering, University of California, Berkeley, California, USA; Environmental Genomics and Systems Biology Division, Lawrence Berkeley National Laboratory, Berkeley, California, USA

**Keywords:** Genome-scale metabolic model, Flux balance analysis, RB-TnSeq

## Abstract

The *E. coli* genome-scale metabolic model (GEM) is a gold standard for the simulation of cellular metabolism. Experimental validation of model predictions is essential to pinpoint model uncertainty and ensure continued development of accurate models. Here we assessed the accuracy of the *E. coli* GEM using published mutant fitness data for the growth of gene knockout mutants across thousands of genes and 25 different carbon sources. We explored the progress of the *E. coli* GEM versions over time and further investigated errors in the latest version of the model (iML1515). We observed that model size is increasing while prediction accuracy is decreasing. We identified several adjustments that improve model accuracy – the addition of vitamins/cofactors and re-assignment of reaction reversibility and isoenzyme gene to reaction mapping. Furthermore, we applied a machine learning approach which identified hydrogen ion exchange and central metabolism branch points as important determinants of model accuracy. Continued integration of experimental data to validate GEMs will improve predictive modeling of the mapping from genotype to metabolic phenotype in *E. coli* and beyond.

**Synopsis:** 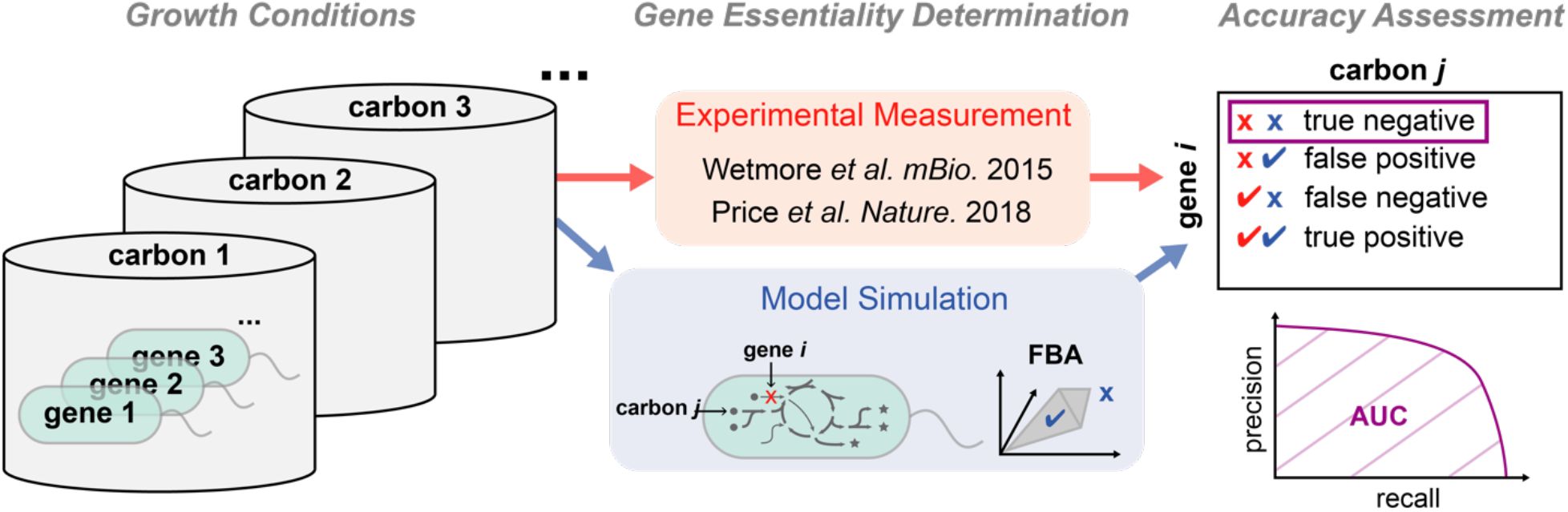

*E. coli* genome-scale metabolic model flux balance analysis (FBA) prediction accuracy was quantified with published experimental data assaying gene knockout mutant growth across different carbon sources. Insights into model development trends and sources of inaccuracy were revealed.

- Model representational power (size) has been increasing over time, while accuracy has been decreasing.
- Adding vitamins/cofactors to the model environment and re-assigning reaction reversibility and isoenzyme gene-to-reaction mapping improves correspondence between model predictions and experimental data.
- Machine learning reveals hydrogen ion exchange and central metabolism branch points as important features in the determination of model accuracy.

## Introduction

The *E. coli* genome-scale metabolic model (GEM) represents one of the most well-established compendia of knowledge on a single organism’s cellular metabolism. This model maps genotype to metabolic phenotype and can be used to mechanistically simulate *E. coli* growth under various gene knockouts and/or environmental chemical perturbations. The *E. coli* GEM was one of the first GEMs to be analyzed (Varma and Palsson 1994), and has undergone iterative curation for over 20 years (Reed et al. 2003; Feist et al. 2007; Orth et al. 2011; Monk et al. 2017). The *E. coli* GEM serves as a gold standard both for the reconstruction of new GEMs for other organisms and for benchmarking our ability to quantitatively simulate metabolism at the genome-scale (Machado et al. 2018; Zimmermann, Kaleta, and Waschina 2021; Henry et al. 2010).

Despite success in mapping the *E. coli* genome to metabolic functions, uncertainty in GEM reconstruction and analysis still generally limits our ability to accurately simulate metabolic phenotypes (Bernstein et al. 2021). For example, specifications of gene-to-reaction mappings, or the chemical composition of the environment for specific experiments can differ from researcher to researcher or computational pipeline to pipeline (Mendoza et al. 2019). Furthermore, it is not always clear how to optimally simulate metabolic flux in the cell given regulatory and other non-metabolic constraints. As we continue to reconstruct GEMs for new organisms these issues are more prominent (Ankrah et al. 2021).

Critical assessment of model prediction accuracy, using experimental data, is essential for pinpointing sources of model uncertainty and ensuring continued development of accurate models. One rich source of data that can be used to validate GEMs is high-throughput mutant phenotype measurements – as measured through random barcode transposon-site sequencing (RB-TnSeq) (Wetmore et al. 2015; Price et al. 2018). This approach utilizes the power of highly parallelized genetic library screens to assay the fitness of gene knockout mutants across an array of conditions. The data that is generated can be readily simulated by GEMs and has been used recently to curate metabolic models (diCenzo, Mengoni, and Fondi 2019; Ong et al. 2020), and benchmark several new automated GEM reconstruction pipelines (Machado et al. 2018; Zimmermann, Kaleta, and Waschina 2021).

In this work, we provide a critical assessment of the *E. coli* genome-scale metabolic model’s accuracy using high-throughput mutant phenotype data measuring the fitness of *E. coli* gene knockout mutants for thousands genes grown across environments containing 25 different primary carbon sources (Wetmore et al. 2015; Price et al. 2018). We compare the size and accuracy of the four latest *E. coli* GEMs to outline progress in the field (Reed et al. 2003; Feist et al. 2007; Orth et al. 2011; Monk et al. 2017). We then perform a detailed investigation of the errors in the latest *E. coli* GEM (iML1515). We identify straightforward adjustments to the model that can improve accuracy and use a machine learning framework to suggest specific fluxes associated with incorrect model predictions.

## Results

### Progression of *E. coli* Genome-Scale Metabolic Models

We calculated the accuracy of the *E. coli* GEM by comparing model predictions to previously published experimental data (Wetmore et al. 2015; Price et al. 2018). We generated model predictions for each experiment by knocking out the specified gene and adding the specified carbon source to the simulation environment and simulating a growth/no-growth phenotype with flux balance analysis (FBA). We then quantified the accuracy of the model based on the area under a precision recall curve (AUC) (Figure 1A, B; see methods for additional details). The precision and recall calculations for this metric focused on true negatives (defined as experiments with low fitness and model predicted gene essentiality). This metric was chosen, as opposed to the overall accuracy or receiver operating characteristic, because the highly imbalanced nature of the data set (far more positives than negatives; Figure 1A inset) suggests that the correct prediction of gene essentiality is more biologically meaningful than the converse prediction of gene non-essentiality.

**Figure 1:**
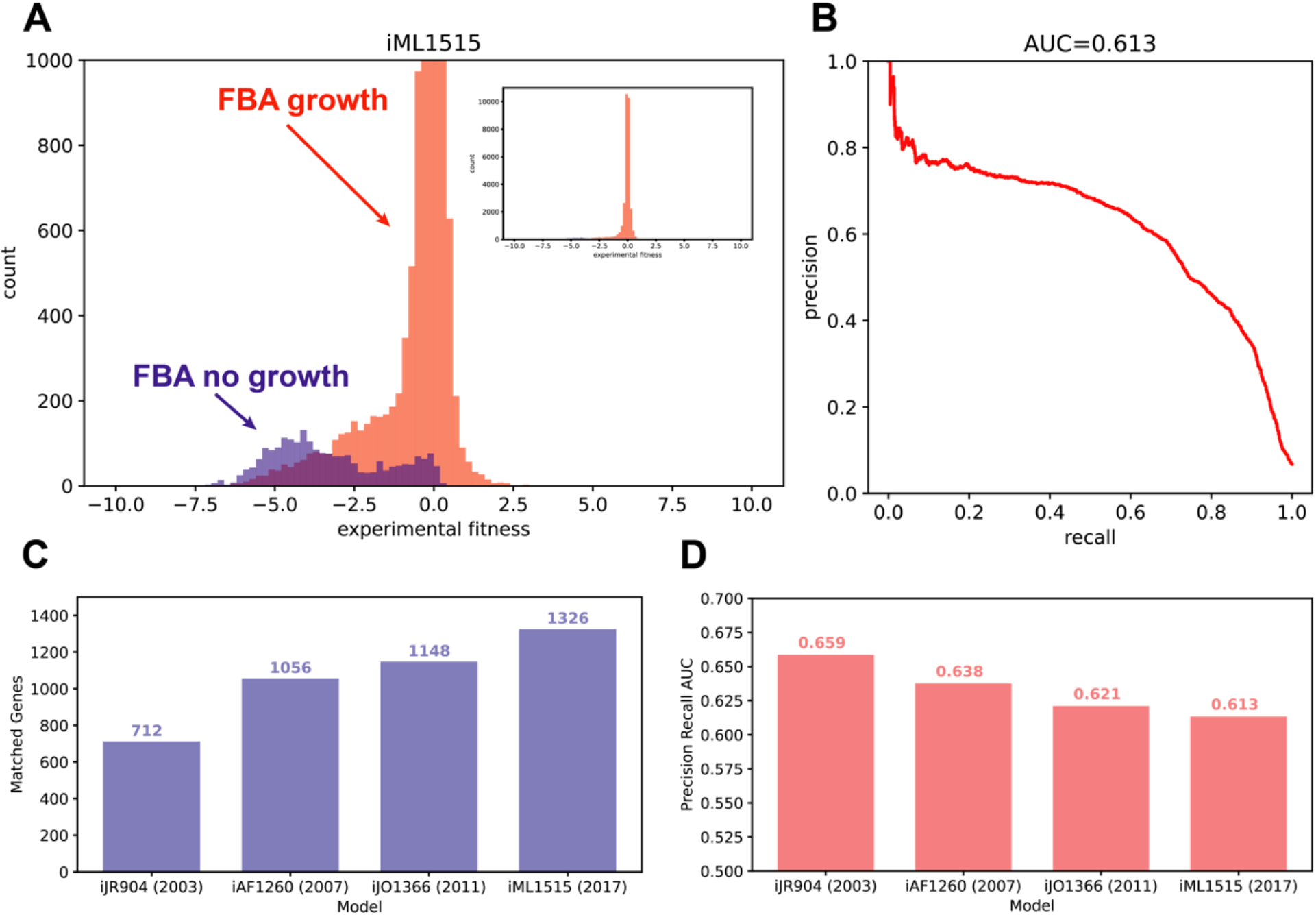
Comparison of *E. coli* GEM accuracy for four subsequent versions of the model. **A)** A histograms of model predictions and experimental fitness data are used to visualize the accuracy of the model. Predictions with flux balance analysis (FBA) biomass flux < 0.001 (no-growth) are included in the blue histogram, and >= 0.001 (growth) in the red histogram. The results for the iML1515 model are shown here. The histogram is cut off at 1000 counts, and the inset (cutoff at 10000 counts) shows the full histogram. **B)** The area under a precision recall curve (AUC) is used to quantify model prediction accuracy. The precision recall curve is calculated using the fitness value as a threshold to predict model essentiality. The iML1515 curve is shown. **C)** The number of genes matched between the model and the experimental data set across subsequent *E. coli* GEMs is shown. **D)** The accuracy of the models is shown across subsequent *E. coli* GEMs, as measured by area under precision recall curve.

We began by comparing the accuracy of four versions of the *E. coli* GEM, which have been subsequently curated from 2003-2017 (iJR904, iAF1260, iJO1366, and iML1515)(Reed et al. 2003; Feist et al. 2007; Orth et al. 2011; Monk et al. 2017). We observed that the number of genes matched between the model and the data set has steadily increased (Figure 2 C). This indicates the increasing power of genome-scale metabolic models to capture metabolic functions. However, the accuracy of the models, as measured by the AUC, has steadily decreased (Figure 2 D). This slight decrease in accuracy points to the importance of balancing increases in model representational power with benchmarks of model accuracy.

**Figure 2:**
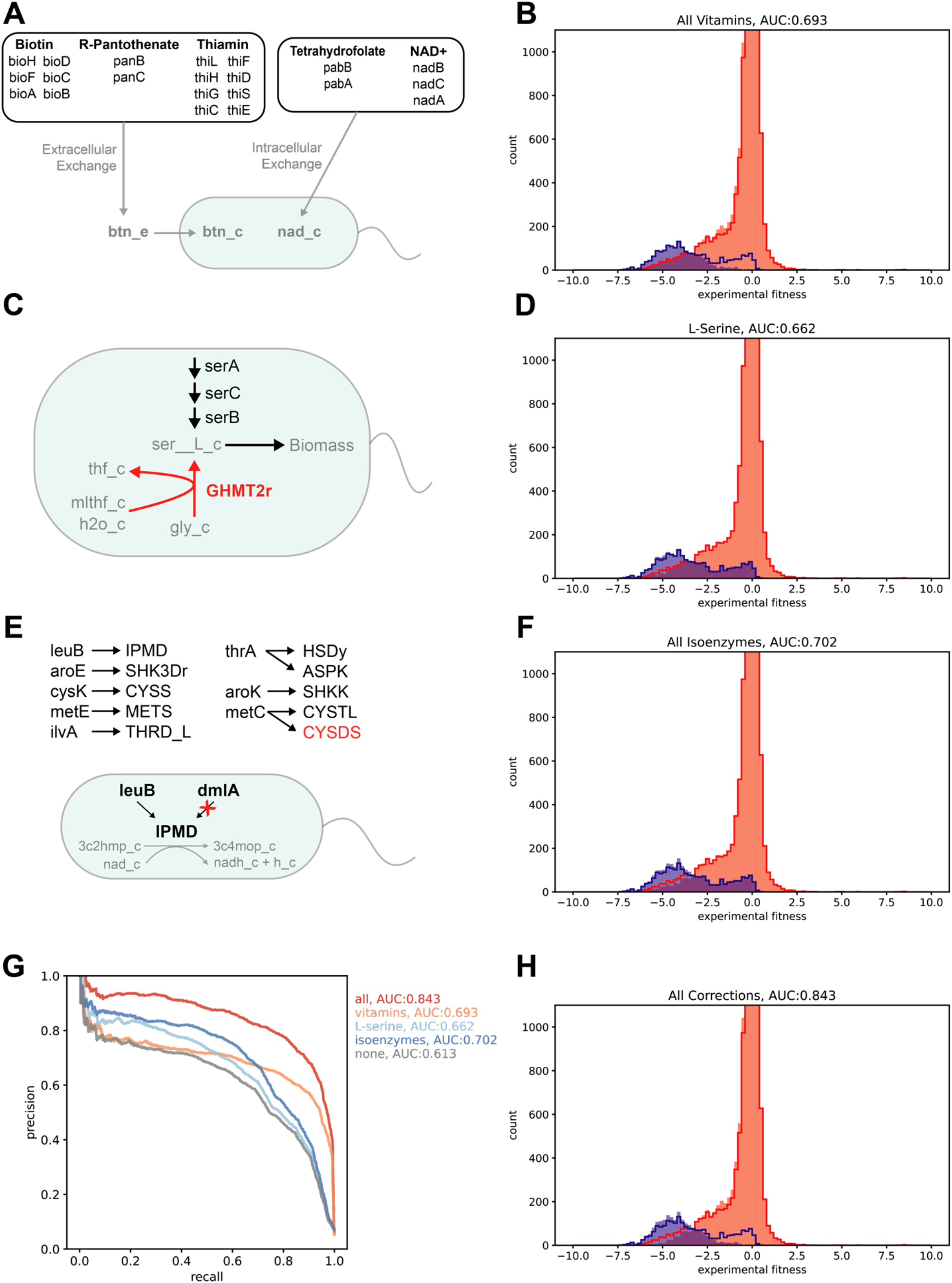
Correction of errors in the latest *E. coli* GEM (iML1515) **A)** Adding vitamins and cofactors to the model environment corrected false negatives (high fitness, model essential). Several genes in vitamin/cofactor biosynthesis pathways were among the highest average fitness with model predicted essentiality across all carbon sources. These genes are listed, grouped by their biosynthetic pathway. Vitamins/cofactors are further grouped into extracellular exchange and intracellular exchange based on whether the model contained a transporter for the associated extracellular metabolite. These vitamins/cofactors were added to the model extracellular or intracellular space through existing or newly added exchange reactions. **B)** The model prediction accuracy, with vitamins/cofactors added, is displayed as the fitness histogram. The area under the precision recall curve (AUC) is listed in the title. The original histogram, without vitamins/cofactors added, is shown as red and blue outlines. The addition of vitamins/cofactors corrected many false negative predictions, as seen by the decrease in the component of the blue histogram with high fitness values. **C)** L-serine biosynthesis gene essentiality predictions are corrected by adjusting the reversibility of the GHMT2r reaction. Three genes in the L-serine biosynthetic pathway (serA, serC, and serB) had incorrect false positive predictions (low fitness, model non-essential). Negative flux through the GHMT2r reaction creates an alternative route for L-serine biosynthesis. Adjusting this reaction to be irreversible makes the L-serine biosynthetic genes essential and corrects the false positive predictions. **D)** The model prediction accuracy, with the GHMT2r reaction made irreversible, is displayed as the fitness histogram. The area under the precision recall curve (AUC) is listed in the title. The original histogram, with GHMT2r reversible, is shown as red and blue outlines. The adjustment of GHMT2r reversibility corrected false positive predictions as shown by the slight decrease in the red histogram below the red outline for low fitness values. **E)** Adjustment of isoenzyme gene-to-reaction mapping corrected false positive predictions for several genes. Isoenzyme genes, with low fitness and model predicted non-essentiality, are listed with an arrow pointing to the reactions for which they are an isoenzyme. Gene-to-reaction mapping was adjusted to make these isoenzymes solely responsible for their corresponding reactions. An example where the lueB gene is mapped to the IPMD reaction by removing the alternative mapping of the dmlA gene to this reaction is shown. Adjusting the gene-to-reaction mapping improved model prediction accuracy for all isoenzymes, excluding metC to CYSDS which had no impact (shown in red). **F)** The model prediction accuracy, with adjusted isoenzyme gene-to-reaction mapping, is displayed as the fitness histogram. The area under the precision recall curve (AUC) is listed in the title. The original histogram, with original isoenzyme gene-to-reaction mapping, is shown as red and blue outlines. The adjustment of isoenzyme mapping corrected false positive predictions as shown by the slight decrease in the red histogram below the red outline for low fitness values. **G)** The precision recall curve is shown for the original uncorrected model, each of the above corrections, and with all the corrections combined. The area under the precision recall curves is listed in the legend to the right of the figure. **H)** The model prediction accuracy, with all corrections combined, is displayed as the fitness histogram. The area under the precision recall curve (AUC) is listed in the title. The original histogram, with no corrections, is shown as red and blue outlines. The corrections led to a decrease in both false positive and false negative predictions.

### Correction of Errors in the iML1515 Model

We sought to investigate the major sources of errors in the latest *E. coli* GEM (iML1515), and to implement model adjustments that could correct these predictions. Several key areas contributing to poor performance were apparent upon visualization and analysis of the model predictions (Figure 2, Supplemental Figure S1).

First, many genes involved in vitamin and cofactor biosynthesis were leading to false negative predictions (Figure 2A). A total of 21 different genes involved in the biosynthesis of biotin, R-pantothenate, thiamin, tetrahydrofolate, and NAD+ were implicated. These genes, when knocked out of the model, create a growth defect. However, the experimental fitness of the corresponding gene knockouts was high. These predictions were corrected by adding the vitamins/cofactors to the simulation environment. They were either added to the extracellular (extracellular exchange) or cytoplasmic (intracellular exchange) compartment of the model, depending on if the model already contained a transporter and exchange reaction for the vitamin/cofactor. Addition of each individual vitamin/cofactor improved model accuracy, and addition of all led to substantial improvement in accuracy (Figure 2B). This result indicates that the identified vitamins/cofactors may be available to the mutants in the RB-TnSeq experiments.

Possible mechanisms for the availability of vitamins/cofactors in the experiments include cross-feeding between the diverse library of *E. coli* mutants or carry-over within individual *E. coli* mutants. We examined an alternative set of experimental RB-TnSeq data, collected at 5 and 12 generations for *E. coli* grown in a minimal glucose medium (Price et al. 2016). This data showed that the phenotypes for genes in the biosynthetic pathways of R-pantothenate (panB, C), thiamin (thiC-H), and NAD+ (nadA-C) had weak negative fitness after 5 generations, but fitness dropped off to be strongly negative after 12 generations. This pattern supports the carry-over hypothesis and suggests that increasing the number of experimental generations could correct these false negative predictions. Alternatively, genes in the biosynthetic pathways for biotin (bioA-D, F, H) and tetrahydrofolate (pabA, B) showed weak negative fitness at both 5 and 12 generations. After 12 generations these metabolites would be depleted by around a factor of 2^12^ (>1000x). This suggests that carry-over alone could not maintain the observed growth in these mutants. A separate study – using the Keio collection of individual gene knockout mutants across 30 carbon sources – reported that knockouts of these genes in the biotin and tetrahydrofolate pathways were not essential when assayed on solid medium (where diverse neighboring colonies could in principle cross-feed metabolites) but were essential when grown in individual liquid cultures (Tong et al. 2020). This suggests that biotin and tetrahydrofolate (or precursors of these metabolites) are cross-fed between *E. coli* mutants. Further in line with the carry-over and cross-feeding hypotheses, it has been demonstrated that many vitamin/cofactor precursors are stable and persist for several generations in *E. coli* and other organisms, and that diverse auxotrophs’ growth is supported by co-culture with prototrophs (Hartl et al. 2017; Ryback, Bortfeld-Miller, and Vorholt 2022). Considering this evidence, the cross-feeding and carry-over hypotheses should be considered when assessing the accuracy of GEM reconstruction pipelines and implementing gap-filling approaches using high-throughput mutant phenotyping data. For example, if these metabolites are present in the experiments but not added to the simulation environment it could lead to the addition of new gap-filled biosynthetic reactions that introduce false positive predictions in more well controlled environments.

Next, we focused on genes that were contributing to false positive predictions. These did not cause a growth defect when knocked out of the model but had low experimental fitness values. Three genes in the L-serine biosynthesis pathway (serA, serB, and serC) were implicated (Figure 2C). Through examination of the metabolic flux in the gene knockout models it was observed that L-serine was being produced from glycine by a reversible reaction (GHMT2r, glycine hydroxymethyltransferase). Setting this reaction to be irreversible corrected the essentiality predictions for the three genes in the L-serine biosynthetic pathway and improved the overall accuracy of the model (Figure 2D). This correction points to reaction reversibility as a key variable in curating GEMs to match experimental data. The reaction in question here has been proposed to run in reverse as a possible route for L-serine biosynthesis from glycine. However, the function of this reaction for this purpose is not firmly established (Price, Deutschbauer, and Arkin 2020).

Another set of genes that were observed to contribute to false positive predictions were genes involved in isoenzyme gene-to-reaction mappings (where there is an “or” relationship in the Boolean mapping of genes to a reaction). Eight different isoenzymes mapping to ten different reactions were among the lowest fitness genes for which the model simulated a growth phenotype (Figure 2E). Reassigning the gene-to-reaction mapping for each of these isoenzyme/reaction pairs, such that each gene was solely responsible for the reaction, improved model performance for all but one pair (metC, CYSDS). The metC gene is mapped to two reactions as an isoenzyme (CYSDS and CYSTL). Only CYSTL is essential in minimal carbon medium. Adjusting the isoenzyme mapping for the essential reaction (CYSTL) improved model accuracy but adjusting the mapping for the CYSDS had no effect on accuracy. Reassigning all isoenzyme gene-to-reaction mappings (excluding metC, CYSDS) led to a further increase in model accuracy (Figure 2F). This correction suggests that isoenzyme representation is an important area for continued curation of GEMs. Isoenzymes can be difficult to properly account for in metabolic models, as different enzymes may be expressed under different regulatory states (Jacobs et al. 2017; Ihmels, Levy, and Barkai 2004). Thus, representations of isoenzymes that do not account for regulatory information can generate overly promiscuous metabolic networks leading to false positive predictions. One example is shown in Figure 4C where dmlA can replace the function of leuB. However, dmlA expression is induced by the presence of D-malate (greater than 50-fold relative to expression in L-malate, D-glucose, or glycerol) (Stern and Hegre 1966). Thus, dmlA would not rescue leuB mutants in many conditions.

Altogether, the three corrections mentioned above, none of which is carbon source specific, substantially improved overall model prediction accuracy (Figure 2G, H, Supplemental Figure S2). These corrections nearly eliminated false negative predictions and substantially reduced false positive predictions. Importantly, they point to several specific areas of GEM reconstruction where adjustments can be made to improve correspondence between model predictions and experimental data.

### Carbon Source Specific Predictions

The dataset utilized here assays 25 different carbon sources, providing insight across diverse carbon utilization pathways. We explored the carbon source specific accuracy of the corrected iML1515 model by calculating the area under the precision recall curve for predicting gene knockout phenotypes for each separate carbon source (Figure 3A). We observed that gluconeogenic carbon sources and other carbon sources that enter metabolism below glycolysis appeared to have lower accuracy than glycolytic carbon sources. This may indicate that our representation of carbon source utilization pathways is more accurate for glycolytic substrates than for other alternative pathways. Additionally, we observed that carbon source specific gene knockout model essentiality predictions were more likely to occur in genes coding for reactions that are near the specified carbon source in the metabolic network (Figure 3B). This should be expected as genes in the pathway for utilization of specific carbon sources are likely to be important for growth on those substrates. It suggests that these genes are the main contributors to carbon source specific predictions rather than global metabolic processes.

**Figure 3:**
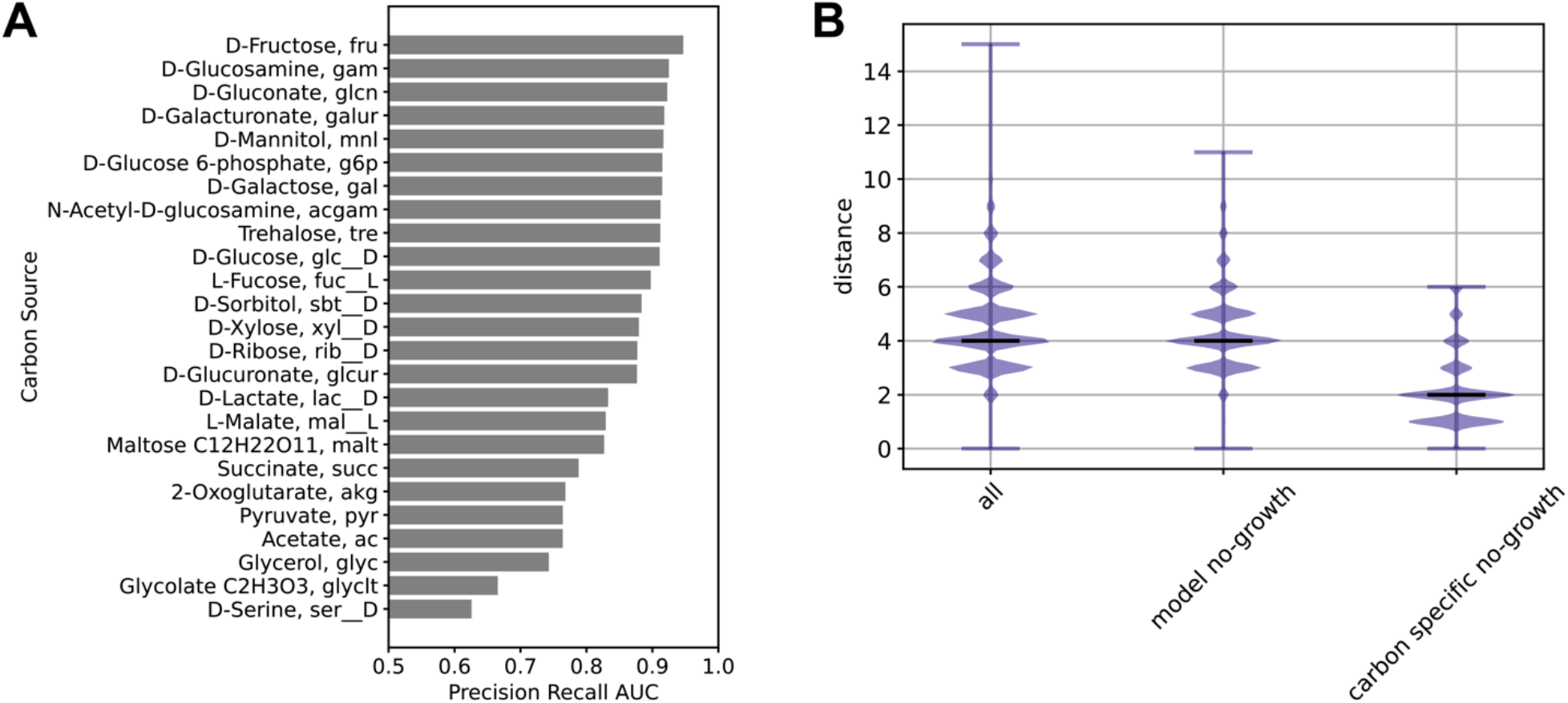
Carbon source specific metabolic model predictions. **A)** The accuracies of the model predictions (area under the precision recall curve, with all corrections) for each specific carbon source are shown. **B)** The distance along the metabolic network between the carbon source and knocked out gene is shown for different subsets of experiments. The distance is calculated as the number of reactions from carbon source to the closest reaction for which the gene is essential (see methods for additional details). The distance is plotted as a violin plot with short bandwidth filter to show the distribution of distances at each integer value. The “all” subset of data shows the distribution of distances for all experiments involving gene knockouts that disrupt at least one reaction. The “model no-growth” subset shows the distribution of experiments with simulated no-growth. The “carbon specific no-growth” subset shows the distribution of experiments where there was simulated no-growth that was carbon source specific (no growth in 80% of carbon sources or less).

### Machine Learning Suggests Flux Profiles Associated with Incorrect Predictions

Genome-scale metabolic modeling provides additional insight beyond a prediction of growth/no-growth. Each simulation, where a growth phenotype is predicted, simultaneously predicts the metabolic flux through every reaction in the network. We sought to use this flux information to gain deeper insight into model accuracy. We began by calculating the metabolic fluxes for each simulation using parsimonious flux balance analysis (Lewis et al. 2010). Visualization of the flux space for each simulation, through principal component analysis, revealed centers for each carbon source wild-type flux distribution surrounded by clouds of the gene knockout simulations grown with that carbon source (Figure 4A). The Euclidean distance of the knockout flux vector from the wild-type flux had a slight negative correlation with the experimental fitness value (Figure 4B). This indicated a weak relationship suggesting that gene knockouts that perturb wild-type flux more greatly led to larger fitness defects.

**Figure 4:**
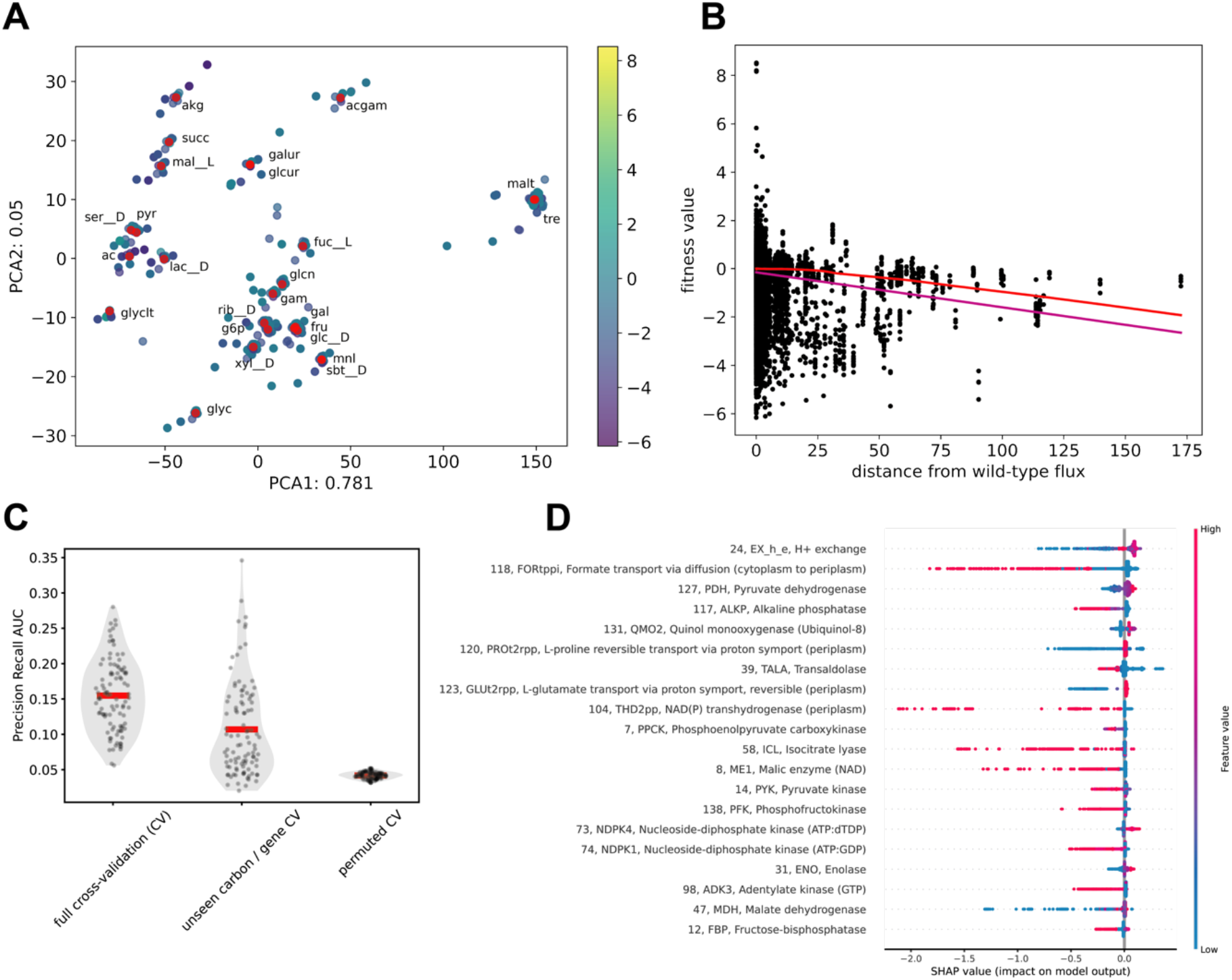
Machine learning with metabolic fluxes to investigate *E. coli* GEM (iML1515) false positive predictions. **A)** The principal component analysis plot of experiments grouped by parsimonious flux balance analysis simulated metabolic flux is shown. Only experiments with simulated biomass flux > 0.001 are included. Wild-type carbon source fluxes (w/ no gene knockouts) are shown as red points. Experiment points are colored according to experimental fitness value. **B)** The plot of experimental fitness value as a function of the Euclidean distance of simulated fluxes from the wild-type fluxes on the same carbon source is shown. A slight negative correlation is seen between fitness and distance (Pearson ρ = −0.175). The purple line shows a linear fit to the data, and the red line shows a lowess fit. **C)** The cross validated accuracy of the machine learning algorithm using simulated fluxes to classify experiments as false positives (fitness values < −2) or true positives (fitness values >= −2) is shown. Only experiments with simulated growth are used here since the simulated fluxes are used as the input for the machine learning algorithm. Precision recall area under the curve is calculated for prediction of false positives. The training set consists of data points from a random subset of 80% of carbon sources and 80% of genes (64% of samples). The full cross-validation (CV) test set contains all remaining data points (36% of samples). The unseen carbon/gene cross-validation set consists of data points that do not involve any of the carbon sources or genes in the training set (4% of samples). The permuted crossvalidation shows prediction accuracy with experimental fitness values for the test set randomly permuted. Violin/scatter plots and means (red line) are shown for 100 random train/test splits. **D)** The feature importance for prediction of data points as true positives calculated through SHAP is shown. Features are sorted by mean absolute SHAP value. For each feature, each sample is shown as a colored point, with the SHAP value on the x-axis and the feature value displayed through the color. When high feature value correlates with high SHAP value, this indicates that high flux through this feature is associated with correct model predictions (true positives). Inversely, high feature values correlating with low SHAP values indicates that high flux through this feature is associated with incorrect model predictions (false positives). SHAP values are averaged across 100 random train/test splits. Feature reaction index, identifier, and name are shown. Each feature corresponds to a representative flux from a flux cluster (see methods for additional information).

To gain additional insight into which flux profiles are contributing most to the accuracy of the model, we used a machine learning approach. We used a gradient boosting decision tree framework, lightGBM (Ke et al. 2017), to classify experiments as true positives or false positives based on their simulated flux profile. Our analysis here focused on positive predictions, as negative predictions (model simulated no-growth) do not have corresponding fluxes to use as the input for the model. The model accuracy was assessed for repeated train/test splits with cross-validation on all experiments (carbon source and gene combinations) that were held out of the training set, and on an orthogonal test set consisting only of experiments with new carbon sources and genes that were not used in the training set (Figure 4C, Supplemental Figure S3). The orthogonal test set provides a measure of the model’s ability to capture metabolic processes that generalize to unseen carbon sources and genes. While this machine learning approach had weak performance, it was able to classify samples better than random, and capture general metabolic processes (Figure 4C).

Next, we utilized Shapley additive explanations, SHAP values (Lundberg and Lee 2017; Lundberg et al. 2020), to quantify the importance of different fluxes in the machine learning (Figure 4D). This analysis revealed flux distributions that were associated with correct or incorrect predictions of the model. Several notable patterns are highlighted (Supplemental Figure S4). The most important feature was the flux of hydrogen ions into or out of the cell. The machine learning model suggested that a hydrogen ion exchange flux close to 0 was associated through SHAP with correct model predictions, a large positive hydrogen ion exchange (ions leaving the cell) was associated with incorrect model predictions, and a large negative hydrogen ion flux (ions entering the cell) was associated strongly with incorrect model predictions (Supplemental Figure S4 A). Several of the other most informative features were also involved in hydrogen ion transfer between the periplasmic and cytoplasmic compartments of the model. The NAD(P) transhydrogenase (THD2pp) uses a flux of hydrogen ions from the periplasm to cytoplasm to reduce NADP+ to NADPH using NADH. High flux through THD2pp was associated with incorrect model predictions. Two symporter reactions (PROt2rpp and GLUt2rpp) transport either L-proline or L-glutamate from the periplasm to cytoplasm along with a hydrogen ion. High negative flux through these reactions (transporting amino acids and hydrogen ions from the cytoplasm to the periplasm) was associated with incorrect model predictions. The flux through both of these reactions was also clustered (strongly covaried across simulations) with a sodium ion symporter that carried the opposite flux transporting the amino acid and a sodium ion back into the cytoplasm. Thus, the net flux of these reaction clusters is the export of hydrogen ions from cytoplasm to periplasm and import of sodium ions. To further address the hydrogen ion flux we re-simulated flux balance analysis growth predictions while fixing the hydrogen ion flux (ranging between 0 and 10) (Supplemental Figure S5). Fixing the hydrogen ion flux to a small positive value increased model accuracy by making genes in the succinate dehydrogenase complex (sdhA-D) essential on acetate and genes in the cytochrome bo complex (cyoA-D) essential on glycolate. Further constraining the hydrogen ion flux to higher values sharply decreased model accuracy by introducing false negative predictions. Beyond the corrections identified by tuning the hydrogen ion flux, there may be more fundamental corrections to GEMs that can be implemented through a more careful representation of hydrogen ion fluxes and cross-membrane gradients.

There were also several reactions involved at branch points in central carbon metabolism that were implicated in the SHAP feature importance analysis. Increased flux through pyruvate dehydrogenase, directing pyruvate to the TCA cycle, was associated with correct model predictions (Supplemental Figure S4 B). Increased flux of glyceraldehyde 3-phosphate through lower glycolysis was associated with correct predictions, while negative flux through lower glycolysis was associated with incorrect predictions (Supplemental Figure S4 C). This result is in line with the previous observation that gluconeogenic carbon sources had lower accuracy. Alternatively, increased flux of glyceraldehyde 3-phosphate through transaldolase in the pentose phosphate pathway was associated with incorrect predictions (Supplemental Figure S4 D). All together, these results suggest that hydrogen ion flux, as well as several major branch points in central carbon metabolism are global determinants of model prediction accuracy.

## Discussion

In this work, we used high-throughput mutant fitness data to conduct a comprehensive analysis of the accuracy of the *E. coli* GEM. The *E. coli* GEM is a gold standard for metabolic model curation, and the dataset we utilize for validation is one of the largest consistent data sets quantifying microbial phenotypes available. We identify several adjustments that can be made to the latest iML1515 metabolic model to improve prediction accuracy. These adjustments highlight including vitamins/cofactors in the simulation environment, and curating reaction reversibility and isoenzyme gene-to-reaction mapping as important areas of uncertainty in GEM reconstruction and simulation. Furthermore, our carbon source specific and machine learning results point to gluconeogenic carbon sources, hydrogen ion flux, and several key branch points in central metabolism as areas of metabolism where GEM predictions warrant further scrutiny.

It is important to note that the corrections we implement in our analysis are not necessarily the only model adjustments that could correct the false model predictions we have identified. In the addition of vitamins/cofactors to the model simulation environment, it is possible that alternative precursors, rather than the vitamins/cofactors themselves, may be the metabolites being cross-fed or carried-over. For example, because the only genes implicated in the tetrahydrofolate analysis were pabA and pabB it is likely that an upstream precursor such as 4-aminobenzoate (PABA) is cross-fed rather than tetrahydrofolate. Next, while we believe the vitamin/cofactor predictions can be explained by the cross-feeding and carry-over hypotheses, other examples of false negative predictions may alternatively be due to missing or unknown biosynthetic reactions that need to be gap-filled. For the false positive predictions, corrected by re-assignment of reaction reversibility and isoenzyme mapping, it is possible that corrections to alternative reactions further up in the pathways of interest could correct these errors. Moving forward, it will be important to establish a systematic and quantitative method for scoring the likelihood of different model corrections based on correspondence with experimental data, prior knowledge from the literature, and parsimony. Such a method could be implemented in a Bayesian framework to formalize the reconstruction/curation of GEMs. Furthermore, curation of GEMs with experimental data and literature evidence should embrace a “deep curation” approach where different sources of evidence that converge to inform a particular model parameter are cross-validated against each other (Macklin et al. 2020). Our work sets the stage for the further development of such systematic and automated methods to reconstruct and curate GEMs based on experimental data and suggests several areas where such approaches could be fruitfully applied.

Our finding that *E. coli* GEM size has been increasing over time while accuracy has been decreasing points towards the importance of the metrics used to assess progress in GEM curation. Model accuracy has been addressed in past curations of GEMs, including for *E. coli.* The original iML1515 publication assessed model accuracy using a similar gene essentiality data set to the one used in this work. In this previous work, the authors measured the growth of the Keio collection of *E. coli* mutants across 16 different carbon sources and compared the results to iJO1366 and iML1515 model predictions (Monk et al. 2017). The reported overall accuracies (iML1515 accuracy: 93.4%, iJO1366 accuracy: 89.9%) are close to the overall accuracy that we calculate from our dataset when we set an experimental growth/no-growth threshold of fitness to −2 (iML1515 accuracy: 93.8%, iJO1366 accuracy: 92.8%). This overall accuracy metric does show an improvement from the iJO1366 model to the iML1515 model. However, we believe that the precision recall AUC metric we used in this work is more biologically meaningful as it emphasizes model accuracy in predicting gene essentiality. For example, the addition of a gene to the model with no metabolic function (that has no impact on the model or experiments) would yield an additional true positive prediction for each carbon source. This non-functional gene would be weighted equally with a gene that is essential for growth across all carbon sources under the overall accuracy metric, while the addition of the non-functional gene would have little impact on the precision-recall AUC metric. All together, we believe that an increased focus on GEM prediction accuracy, and the metrics used for accuracy evaluation, will help enable GEMs to deliver on the promise of predicting genotype from phenotype.

One exciting idea for the development of the field in the direction of improved predictive modeling is the implementation of a community competition centered on critical assessment of microbial phenotype predictions. A community survey of metabolic modeling and microbiome researchers recently highlighted model trust/validation as an important focus for moving the field forward, and proposed such a community initiative (Ankrah et al. 2021). This initiative could serve both to motivate standardized GEM validation and to coordinate data collection and organization. As we move forward, we should also keep in mind that the development of models of cellular physiology that go beyond metabolism will likely be necessary. While GEMs currently offer an appealing balance between predictive power and complexity, expanding models to deal with gene regulation, and other cellular processes has the potential to further improve prediction accuracy for both metabolic and non-metabolic phenotypes (Goldberg et al. 2018). In any case, standardized assessments of model prediction accuracy (with large, curated databases of experimental data) will continue to be essential for the successful application of computational modeling.

## Materials and Methods

All the methods used throughout this analysis are documented in a reproducible Python Jupyter Notebook which is available on GitHub at github.com/dbernste/E_coli_GEM_validation. Metabolic model adjustments and simulations were conducted using COBRApy (Ebrahim et al. 2013). Data analysis, visualization, and machine learning were conducted using: jupyterlab (Kluyver et al. 2016), matplotlib (Hunter 2007), numpy (Harris et al. 2020), pandas (McKinney 2010), scipy (Virtanen et al. 2020)[REF], Scikit-learn (Pedregosa et al. 2011), lightgbm (Ke et al. 2017), and SHAP (Lundberg et al. 2020).

### Data Processing

Experimental RB-TnSeq data was collected from the online fitness browser fit.genomics.lbl.gov (Price et al. 2018). The data from *E. coli* BW25113 was used for this analysis. The fitness values (rather than the t scores) were used to represent the fitness, as preliminary analyses indicated that these scores corresponded more closely with model predictions. Genes were matched to the *E. coli* GEM through their “sysName” which corresponds to their BiGG database identifiers (King et al. 2016). Carbon sources were matched by manually searching the BiGG database for metabolite identifiers, other media components were similarly matched to the BiGG database. The two carbon sources sucrose and mannitol were excluded from this analysis because of known issues in the experimental preparation of their media. The fitness data for sucrose has since been removed from the fitness browser, and data for mannitol has been replaced with corrected experimental data (Price, Deutschbauer, and Arkin 2022). All carbon source experiments were conducted in duplicate in the original data. The fitness scores for these duplicates were averaged for comparison to metabolic model predictions. Preliminary analysis indicated that this averaging slightly improved correspondence between fitness values and model predictions.

### Genome-Scale Metabolic Model Simulation and Adjustment

Metabolic models were downloaded from the BiGG database in sbml format (King et al. 2016). Models were loaded and analyzed using COBRApy (Ebrahim et al. 2013). Models were matched to media and carbon source metabolites and exchange bounds were adjusted to add metabolites to the environment. Exchange lower bounds were set to −1000 [mmol/(gdw*hr)] as a default and −10 [mmol/(gdw*hr)] for all carbon sources. Models were adjusted to account for differences between the experimental BW25113 *E. coli* strain and the model MG1655 strain by removing several genes and their corresponding reactions that are not present in the BW25113 strain (Grenier et al. 2014). Models were further adjusted to remove non-conditionally essential genes from the analysis (genes that were essential when all possible exchanges have lower bound set to −1000). Many of these non-conditionally essential genes were already removed from the fitness data as they do not have reliable phenotype measurements in RB-TnSeq experiments due to low representation in the initial library. Therefore, it is possible that the remaining non-conditionally essential genes, matched between model and dataset, could be false negative predictions. We chose to remove all these genes from our analysis for consistency. Model simulations were conducted with model.slim_optimize, which provided substantial speed improvements when only recording the biomass flux, or with parsimonious FBA through cobra.flux_analysis.pfba to record all simulated fluxes (Lewis et al. 2010). A biomass flux of 0.001 [gdw/(gdw*hr)] was used throughout this analysis as a growth/no-growth cutoff.

### Model Accuracy Calculation

Model accuracy was calculated using the area under a precision recall curve. The model simulated FBA biomass flux data was binarized to a growth/no-growth phenotype based on a cutoff of 0.001 (no-growth < 0.001, growth >= 0.001). A sliding threshold on the fitness value was then used to generate a precision recall curve. The positive class was set to simulated essentiality (no-growth phenotype). The area under this precision recall curve was used to quantify model accuracy. Precision recall curves were calculated using sklearn.metrics.pre_rec and area under the curve was calculated using sklearn.metrics.auc.

### Carbon Source Distance Analysis

The distance between carbon sources and genes was calculated using the genome-scale metabolic model. The model was converted to a bi-partite graph with metabolites and reactions as nodes. An edge was placed between any metabolite and reaction where that metabolite was a reactant or product for that reaction (non-zero stoichiometry), and an adjacency matrix was constructed for the metabolic network. From this adjacency matrix, high degree “hub” nodes were removed. Hub nodes were defined as any node with more than 50 connections and consisted of central metabolites such as hydrogen and water, cofactors such as ATP and NAD+, and the *E. coli* Biomass reactions. Pyruvate was also a hub node but was left in the network due to its role as a carbon source in this analysis. This hub-less network was then used to calculate the carbon source gene distance. The distance was calculated as the number of reactions traversed from the extracellular carbon source to the nearest reaction that was essentially encoded by the gene (a gene knockout removed the reaction). For example, distance of 0 corresponded to an essential reaction directly involving the extracellular carbon source, while distance of 1 corresponded to an essential reaction involving any metabolite connected to a reaction of distance 0. Distances on the metabolic network were calculated using the scipy.csgraph.shortest_path method implementation of Dijkstra’s algorithm.

### Machine Learning and Feature Importance

Machine learning (ML) was conducted to classify simulations with biomass flux into false positives (experimental fitness < −2) or true positives (experimental fitness >= −2). A lightgbm classifier was used. Flux vectors, quantifying the simulated flux across all reactions in the metabolic model (2714 total fluxes), were used as the input for the ML algorithm. Fluxes with variance across samples less than 10^-7^ were discarded form the analysis (leaving 579 total fluxes). Fluxes were clustered into groups of fluxes with covariation greater than 0.99 across samples (leaving 172 flux clusters). One representative flux from each cluster was used for the ML input. ML accuracy was assessed by 100 repeated train/test splits. For each train/test split a random subset of samples from 80% of the carbon sources and 80% of the genes was selected as the training set. Cross-validated accuracy was assessed on the full set of test samples as well as a smaller set of samples with no overlapping carbon sources or genes in the training set. Model performance was explored for varying number of leaves in the ML model, an important tuning parameter for over-fitting. In the final model 5 leaves were used, which balanced full cross-validation with new carbon/gene cross-validation. Other parameters were set to the lightgbm classifier default values (boosting type=‘gbdt’. **num_leaves=5**, max_depth=-1, learning_rate=0.1, n_estimators=100, subsample_for_bin=200000, objective=None, class_weight=None, min_split_gain=0.0, min_child_weight=0.001, min_child_samples=20, subsample=1.0, subsample_freq=0, colsample_bytree=1.0, reg_alpha=0.0, reg_lambda=0.0, random_state=None, n_jobs=None, importance_type=‘gain’). Feature importance was calculated using Shapley additive explanations (SHAP) through shap.TreeExplainer (Lundberg et al. 2020). SHAP values shown in the text are for the classification of samples as true positives and averaged across train/test splits.

## Acknowledgements

We would like to acknowledge helpful discussion and feedback from all members of the Arkin lab. The contributions of DBB were supported by the Consortium for Monitoring, Technology, and Verification under U.S. Department of Energy National Nuclear Security Administration, award number DE-NA0003920. The contributions of APA and MNP by Ecosystems and Networks Integrated with Genes and Molecular Assemblies (ENIGMA; http://enigma.lbl.gov), a Science Focus Area program at Lawrence Berkeley National Laboratory, is based upon work supported by the U.S. Department of Energy, Office of Science, Office of Biological & Environmental Research under contract number DE-AC02-05CH11231.

## Supplemental Figures

**Supplemental Figure S1:**
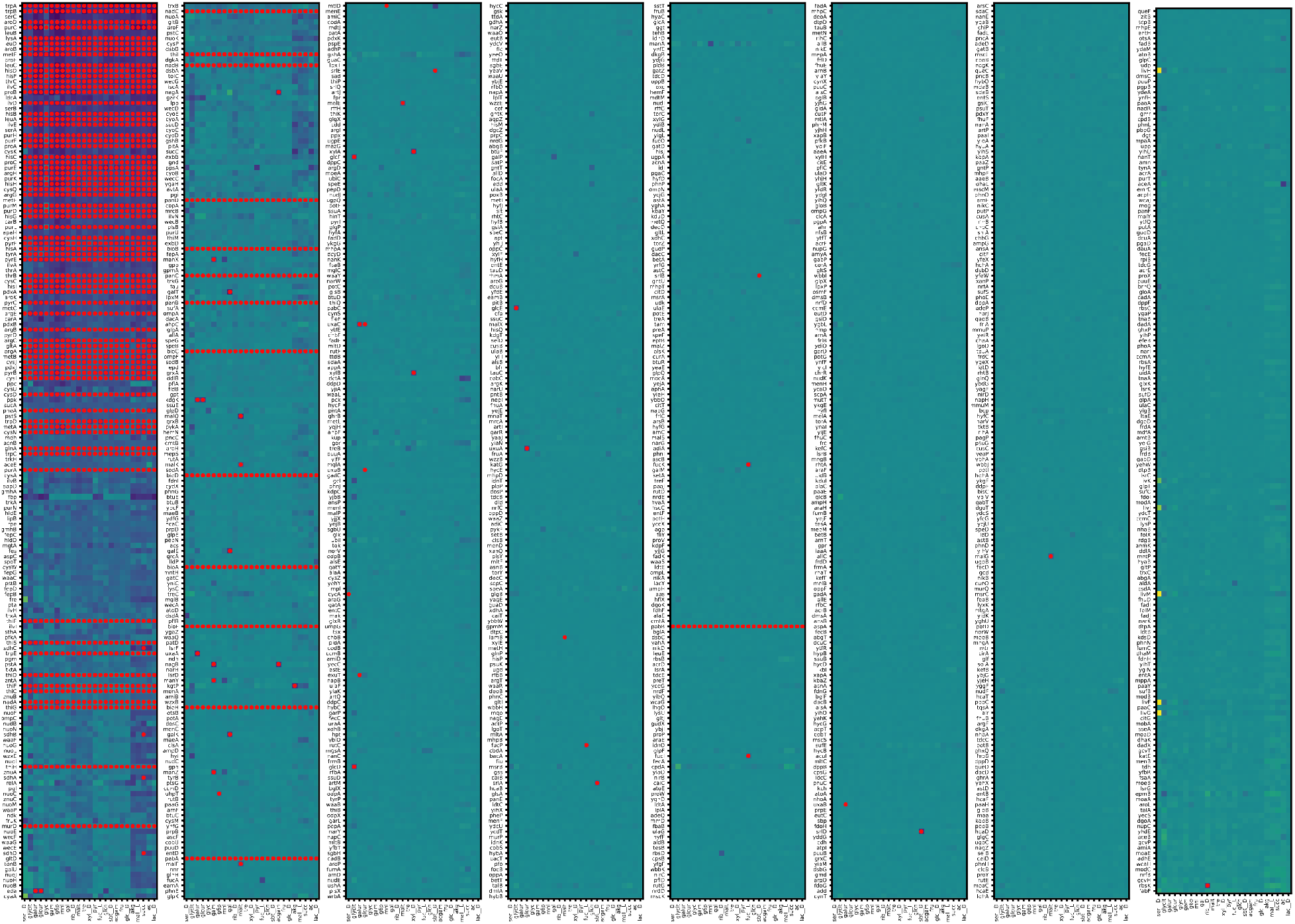
Experimental fitness and simulation results. The entire data matrix of genes by carbon sources is visualized. Color indicates experimental fitness value (dark blue: low, yellow: high), a red dot indicates simulated no-growth (biomass flux < 0.001).

**Supplemental Figure S2:**
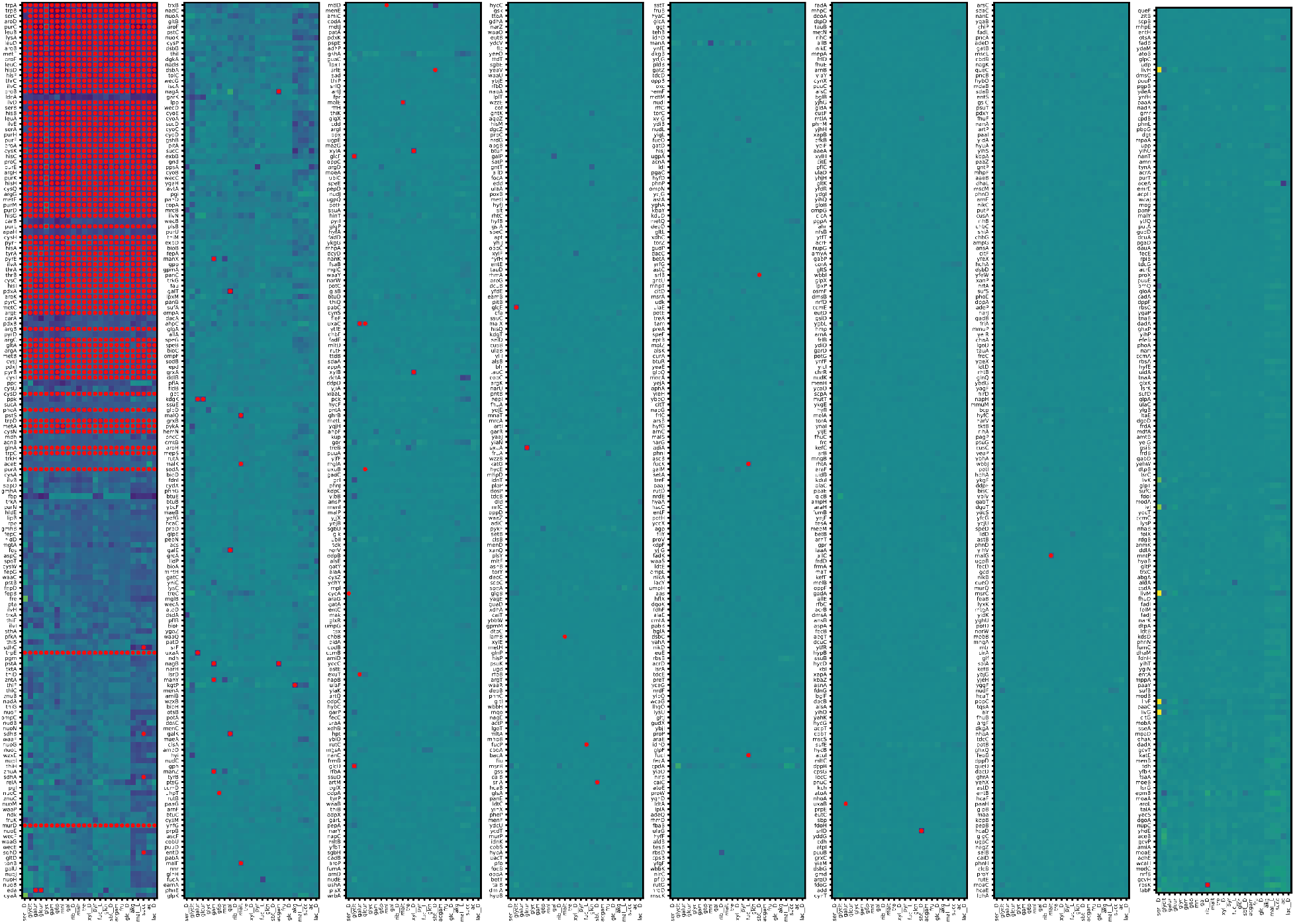
Experimental fitness and post-correction simulation results. The entire data matrix of genes by carbon sources is visualized. Color indicates experimental fitness value (dark blue: low, yellow: high), a red dot indicates simulated no-growth after implementing corrections (biomass flux < 0.001).

**Supplemental Figure S3:**
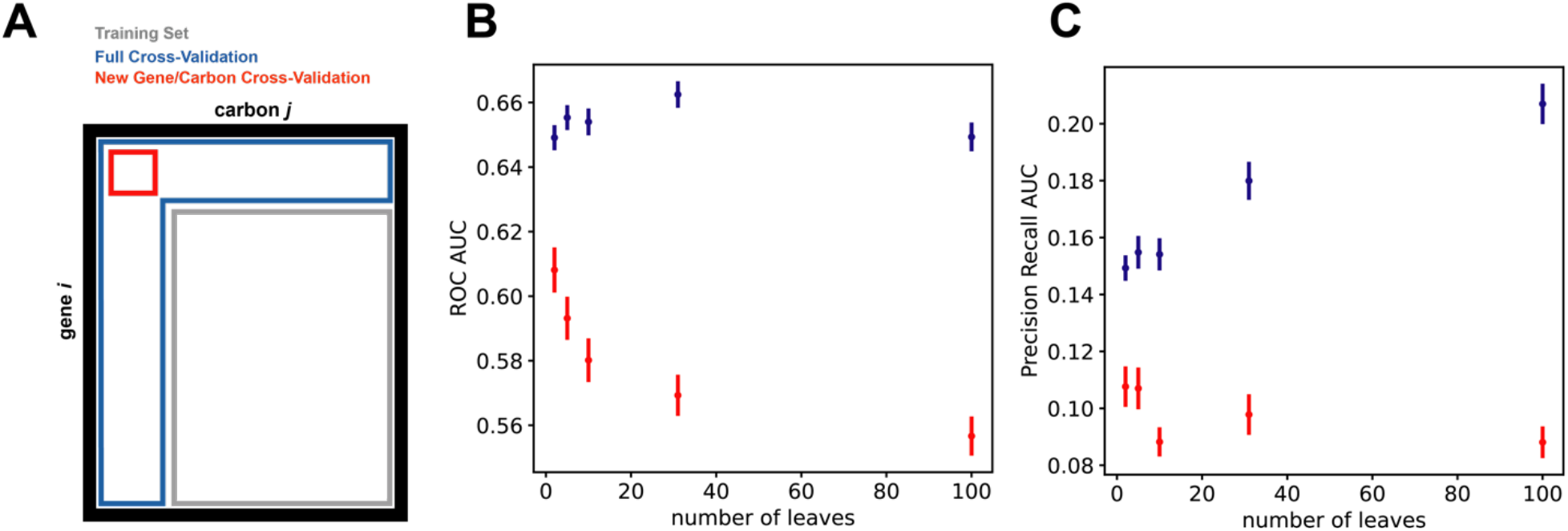
Machine learning training and cross-validation. **A)** The schematic for training and test set partitioning is shown. The data is structured as a matrix of carbon sources and genes (with a vector of metabolic fluxes for each element in this matrix). The test error was calculated in two different ways. 1) For all held out experiments in a full cross-validation (blue outline). 2) For new gene/carbon sources that were unused in the training set (red outline). **B)** The area under the receiver operating curve accuracy of the machine learning model for different numbers of leaves (an important parameter to control overfitting) is plotted for 100 random train/test splits. Mean +/- standard error is shown. **C)** The area under the precision recall curve accuracy of the machine learning model for different numbers of leaves is plotted for 100 random train/test splits. Mean +/- standard error is shown.

**Supplemental Figure S4:**
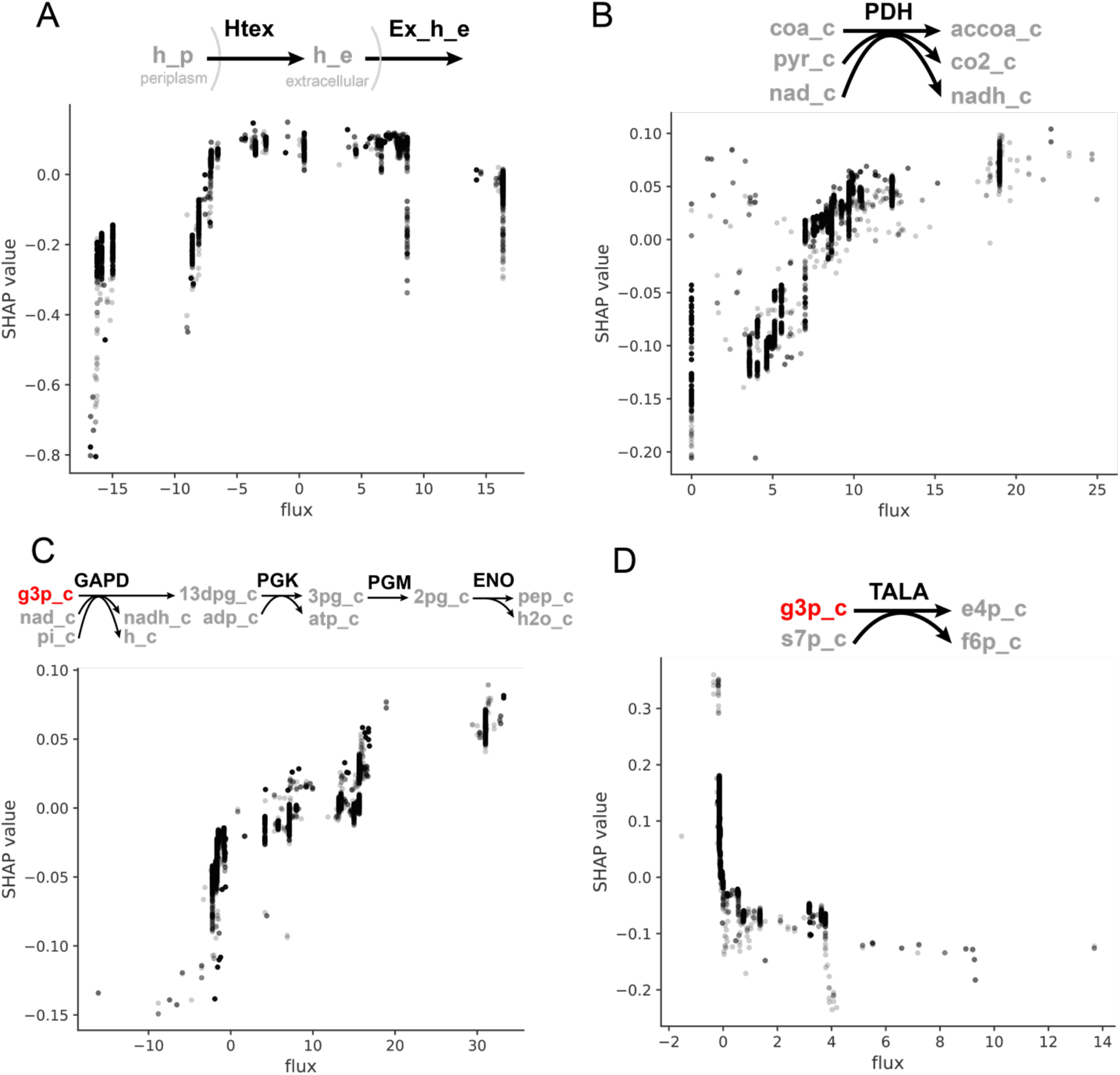
SHAP value dependency plots for select flux features. **A)** Hydrogen ion exchange and transport **B)** Pyruvate dehydrogenase **C)** Lower glycolysis **D)** Transaldolase

**Supplemental Figure S5:**
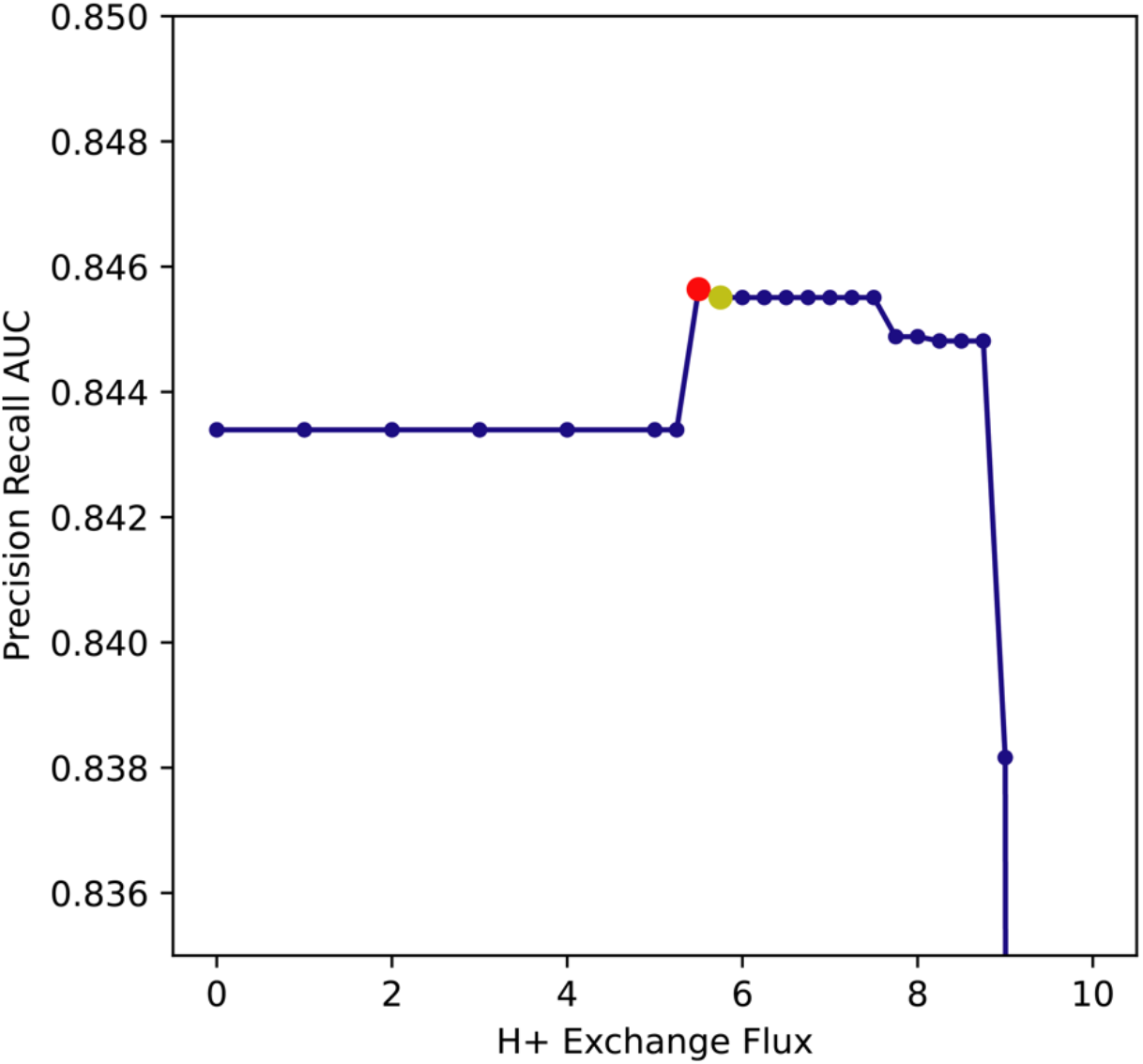
Model prediction accuracy with fixed hydrogen ion flux. Fixing the bounds of hydrogen ion flux to small positive values (dots show tested values) leads to an optimal value that improves model prediction performance. The first uptick in performance comes from the correction of genes in the succinate dehydrogenase complex (sdhA-D) becoming essential for growth on acetate (red dot). The second level of improved prediction performance (which has slightly lower AUC than the red dot) additionally causes genes in the cytochrome complex (CyoA-D) to become essential for growth on glycolate (yellow dot). Further increases in the fixed hydrogen ion flux sharply decrease model prediction performance by adding false negative predictions.

